# A retrospective public external benchmark of healthy-to-stroke lower-limb EEG transport highlights source construction, adaptation burden, and confound sensitivity

**DOI:** 10.64898/2026.03.26.714655

**Authors:** Daniel Choi, Andrew Choi, Quan Lam, Junho Park

## Abstract

**Background:** Lower-limb EEG is a rehabilitation-facing control signal for stroke neurorehabilitation and future non-invasive brain–spine interfaces, but a public external benchmark that jointly audits source construction, minimal adaptation burden, and confound sensitivity is lacking. We therefore tested whether lower-limb effort-versus-rest decoders trained on healthy public EEG transport to a stroke target domain.

**Methods:** We conducted a retrospective public-data external benchmark using three public EEG datasets harmonised to a common lower-limb effort-versus-rest target. Classical and deep models were compared under zero-shot transport, 10-shot temperature calibration, and 10-shot fine-tuning. For few-shot analyses, each target participant contributed a trial-disjoint subject-internal support set of 10 labelled trials per class and a held-out remainder test set. Prespecified analyses audited source construction, support-resampling sensitivity, and montage controls. Uncertainty was summarised with participant-level bootstrap confidence intervals.

**Results:** Within this benchmark, healthy-to-stroke zero-shot transport was weak. The best zero-shot result was classical rather than deep, with CSP+LDA reaching area under the receiver operating characteristic curve (AUROC) 0.603, whereas EEGNet remained near chance (AUROC 0.527). Ten-shot calibration improved operating behaviour more than discrimination: for CSP+LDA, expected calibration error fell from 0.267 to 0.035 and specificity increased from 0.180 to 0.485, whereas AUROC remained essentially unchanged (0.603 to 0.604). Ten-shot fine-tuning produced only modest gains; the best overall AUROC was 0.605 for pooled dataset-balanced CSP+LDA, numerically tied with pooled raw CSP+LDA (0.605). MILimbEEG-only source training was consistently weak, exploratory deep domain-generalisation variants did not rescue transport, and frontal and temporal montage controls remained relatively competitive.

**Conclusions:** Within this public benchmark, source construction and minimal adaptation burden mattered more than model novelty, and retrospective montage controls limited motor-specific interpretation. The results support harmonised prospective validation of lower-limb EEG transport over further retrospective model iteration.

## Background

Recovery of walking and lower-limb function remains one of the most clinically consequential goals after stroke. EEG-based control signals are therefore attractive not only for rehabilitation feedback and assistive interfaces, but also for emerging restorative neurotechnology programmes in which cortical intention could eventually be coupled to peripheral stimulation or spinal neuromodulation.[1–4] Lower-limb EEG is appealing because it is non-invasive, scalable, and already embedded in a literature that spans motor-imagery rehabilitation, gait-intention decoding, and proof-of-concept non-invasive brain–spine interfaces.[1–4]

That clinical promise has outpaced translational validation. Most lower-limb EEG studies are developed and benchmarked within a single dataset, often in healthy participants, and often framed around discrimination performance alone. Yet within-dataset performance does not answer the question that matters for translation: whether a decoder learned in one context survives transfer across subjects, acquisition regimes, and disease states. This problem is not unique to neurotechnology. Across medicine, external validation remains underused, and calibration and model updating are too often treated as secondary considerations rather than primary translational variables.[5–8]

The lower-limb EEG literature is especially vulnerable to this gap. Public EEG datasets differ substantially in acquisition, cueing, label structure, and metadata completeness. A recent review of public motor imagery and execution datasets reported a mean two-class motor-imagery accuracy of 66.53% and an estimated poor-performer fraction of 36.27%, underscoring that even healthy, within-domain EEG decoding is far from uniformly strong.[9] At the same time, clinically relevant lower-limb neurotechnology is moving toward clinical-target applications in stroke rehabilitation and non-invasive brain–spine interfacing, where performance will depend on stroke-domain transport rather than healthy within-dataset accuracy.[2–4]

This question is now testable with public data. EEGMMIDB provides a widely used healthy motor execution and motor imagery resource with more than 1500 recordings from 109 volunteers.[10, 11] MILimbEEG adds lower-limb execution and imagery tasks from 60 participants, but with a markedly different class structure.[12] Stroke2025 provides a public lower-limb EEG dataset involving 27 participants with stroke and 4260 trials, making stroke-domain transport analysable in an open benchmark setting.[13] Together, these resources allow a sharper translational question: if a lower-limb effort-versus-rest decoder is trained on healthy public EEG, what remains when it is transported into stroke? Prior work has already transferred deep-learning motor-imagery models from healthy cohorts to cohorts involving participants with stroke in narrower settings.[14] The novelty claimed here is therefore not the bare idea of healthy-to-stroke transfer itself. It is a fully public lower-limb external benchmark in which source construction, minimal adaptation burden, and confound-sensitive montage controls are audited within one locked evaluation framework.

We designed this study as a translational constraint paper rather than a model-optimisation paper. The objectives were fourfold: first, to quantify healthy-to-stroke zero-shot transport under a leakage-controlled public-data benchmark; second, to determine whether minimal subject-specific adaptation materially improves performance; third, to test whether source construction matters more than model class within this benchmark; and fourth, to assess whether retrospective montage controls support or limit a motor-specific interpretation. The purpose was not to prove a deployable biomarker, but to identify the real bottlenecks that must be cleared before lower-limb EEG control can credibly support neurorehabilitation or future brain–spine interface work.

## Methods

### Study design and reporting frame

We conducted a retrospective external evaluation using public human EEG datasets. The study was designed as a transport benchmark rather than a model-development leaderboard, with the central question being whether lower-limb effort-versus-rest decoders trained on healthy public EEG survive transfer into stroke. The manuscript was structured around principles from TRIPOD+AI for prediction-model evaluation and STROBE for observational reporting.[7, 8] No new participants were recruited.

### Datasets

Three public datasets were used.

1. **EEGMMIDB** (PhysioNet EEG Motor Movement/Imagery Dataset), a healthy motor execution and motor imagery dataset comprising more than 1500 recordings from 109 volunteers.[10, 11] The transport benchmark used a fixed source subset defined in the locked configuration files.
2. **MILimbEEG**, a public lower-limb execution and imagery dataset from 60 participants released with 7440 CSV files.[12]
3. **Stroke2025**, a public lower-limb motor-imagery EEG dataset involving 27 participants with stroke, two enhanced paradigms, three time points, and accompanying patient information.[13]

The benchmark code used fixed YAML configurations for each experiment family within a version-controlled benchmark codebase. Healthy source data were drawn from EEGMMIDB and MILimbEEG; Stroke2025 served as the target transport domain.

Stroke2025 was the sole target domain because it provides public longitudinal stroke EEG rather than a single-session convenience cohort. In the locked benchmark loader, the public group labels PRE, IES, SES, POST, and FOLLOW were preserved as session metadata for downstream splitting and auditing; these labels correspond to a conventional baseline assessment, two stimulation-enhanced initial assessments, a post-treatment conventional assessment, and a follow-up conventional assessment.[13]

### Control variable and label harmonisation

The common control variable was binary lower-limb `effort_rest`. Harmonisation rules were fixed before the transport experiments and summarised in Table 1.

- **EEGMMIDB:** both-feet execution and imagery events (`T2`) were mapped to effort; rest events (`T0`) were mapped to rest. Windows spanned 0.0–2.0 s after event onset.
- **MILimbEEG:** filename class 1 was mapped to rest; classes 4–7 were mapped to effort across execution and imagery modes; classes 2 and 3 were excluded. Trials were resampled or interpolated to a 2.0 s window at 125 Hz.
- **Stroke2025:** event identifiers 3/4/5/6 were mapped to effort and 7/8 to a rest-proxy class in the locked benchmark loader. Windows spanned 0.0–2.0 s after annotation onset. Files lacking both effort and rest-proxy labels were excluded.

**Table 1.** Dataset characteristics, task mapping, and label harmonisation.

| Dataset | Role | Public cohort / subset | Effort mapping | Rest mapping | Windowing | Key exclusions |
| --- | --- | --- | --- | --- | --- | --- |
| EEGMMIDB | Healthy source | 109 volunteers; balanced source subset 908 trials | T2 both-feet execution + imagery | T0 rest | 0.0-2.0 s after event onset at 125 Hz | runs without T0/T2 or zero matched windows |
| MILimbEEG | Healthy source | 60 participants; source subset 2520 trials | File classes 4-7 lower-limb active tasks | File class 1 | Full trial resampled/interpolated to 2.0s at 125 Hz | classes 2/3 excluded from effort rest; class 8 retained only if present |
| Stroke2025 | Stroke target | 27 participants; 4260 trials | Event IDs 3/4/5/6 lower-limb MI | Event IDs 7/8 rest proxy | 0.0s to 2.0s after annotation onset at 125 Hz | files missing both MI and rest-proxy labels excluded |

The harmonised variable was intentionally coarse. EEGMMIDB mixes healthy execution and imagery, MILimbEEG aggregates several lower-limb active classes with different filename semantics, and Stroke2025 is a longitudinal stroke motor-imagery resource. A narrower shared label, such as imagery-only, would have discarded large portions of the already limited public source data while still leaving cueing and acquisition differences unresolved. For Stroke2025 specifically, the locked benchmark used the public 7/8 idle-task annotations as a rest proxy to preserve a consistent binary target across the available longitudinal groups; this should be read as a pragmatic benchmark choice rather than as a phase-matched idle-state physiology contrast. We therefore treated `effort_rest` as the most defensible shared variable for external transport, while explicitly interpreting it as a pragmatic common ontology rather than proof of physiological equivalence across healthy execution, healthy imagery, and stroke motor imagery.

### Preprocessing and windowing

The canonical transport configurations used:

- target sampling frequency 125 Hz,
- band-pass filtering 1–35 Hz,
- average rereferencing,
- per-trial z-scoring,
- fixed 2.0 s windows aligned to the dataset-specific event definitions.

The default deployment-oriented montage for the transport experiments comprised 16 channels: F3, Fz, F4, FC5, FC1, FC2, FC6, C3, Cz, C4, CP5, CP1, CP2, CP6, Pz, and Oz. Physiology-control analyses used alternative fixed montages defined in the synchronized configuration files.

### Montage definitions for the physiology audit

Four montages were compared in the within-dataset EEGMMIDB audit:

- **canonical_16:** F3, Fz, F4, FC5, FC1, FC2, FC6, C3, Cz, C4, CP5, CP1, CP2, CP6, Pz, Oz
- **frontal_control:** Fp1, Fp2, F7, F3, Fz, F4, F8
- **temporal_control:** T3, T4, CP5, CP6, P3, P4, O1, O2
- **motor_laplacian:** FC1, FC2, C3, Cz, C4, CP1, CP2, Pz with surface-Laplacian rereferencing

These controls were intended to limit overinterpretation of the within-dataset signal as narrowly motor-cortical.

### Split strategies

The locked transport benchmark used two split families.

1. **Healthy-to-dataset zero-shot transport:** healthy source data from EEGM-MIDB and MILimbEEG were used for training and validation, and all Stroke2025 data were held as the target test domain. The locked zero-shot configuration used target dataset `stroke2025`, a validation fraction of 0.15 within the healthy source, and fixed random state 42.
2. **Healthy-to-dataset few-shot transport:** support/test independence was enforced at the trial level within participant. For each Stroke2025 participant, all eligible target trials from the included group labels (PRE, IES, SES, POST, FOL- LOW) were pooled across available sessions/time points; 10 labelled trials per class were randomly sampled as the support set, and the non-overlapping remainder from that same participant formed the held-out test set. The main few-shot analyses used fixed support-selection seeds, and targeted robustness analyses later repeated support selection across 20 support draws per participant.

No trial could appear in both support and test. Support and test could nevertheless draw from the same or different public Stroke2025 group labels within a participant, because the locked few-shot benchmark did not enforce session-held-out or timepoint-held-out separation. The few-shot results should therefore be interpreted as subject-internal, trial-disjoint adaptation rather than session-disjoint adaptation.

Source-sampling controls used fixed prespecified seeds and are summarised in Table

### Source-ablation design

Six source conditions were prespecified: EEGMMIDB-only, MILimbEEG-only, pooled raw, pooled class-balanced, pooled dataset-balanced, and pooled class-plus-dataset-balanced. Deterministic source-sampling controls were fixed before analysis. In the raw pooled source, training counts were EEGMMIDB 454 rest / 454 effort and MILimbEEG 120 rest / 2400 effort, for a total of 3428 source trials. Dataset balancing without class balancing reduced MILimbEEG to 41 rest / 867 effort, producing a total of 1816 source trials. Class balancing reduced MILimbEEG to 120 rest / 120 effort while leaving EEGMMIDB unchanged, for 1148 source trials. Full class-plus-dataset balancing yielded 120 rest / 120 effort from each source dataset, for 480 source trials.

### Models

The primary deep model was EEGNet.[15] The primary classical models were:

- CSP + LDA
- Riemannian tangent-space logistic regression (Riemann+TSLR).[16]

An exploratory EEG Transformer comparator implemented in the benchmark codebase was included only in the within-dataset EEGMMIDB physiology audit and is therefore reported only in Table 5 and Figure 5.[17]

Exploratory domain-generalisation variants included CORAL, VREx, GroupDRO, and DANN formulations attached to the EEGNet backbone. Because the paper’s goal was translational evaluation rather than model invention, the deep DG variants were treated as exploratory.

### Training, calibration, and adaptation

Deep models in the transport benchmark used batch size 128, maximum 30 epochs, learning rate 5 *×* 10*^−^*^4^, weight decay 10*^−^*^4^, and patience 6. Fine-tuning used a frozen-backbone configuration with up to 12 epochs, learning rate 10*^−^*^4^, and batch size 16.

Calibration used temperature scaling with threshold selection by balanced accuracy on the designated validation or support set. Temperature scaling fit a single scalar temperature by minimising negative log-likelihood after logit scaling of the model probabilities. After temperature fitting, the operating threshold was selected on the same designated calibration subset by maximising balanced accuracy over a prespecified candidate threshold grid augmented with observed validation probabilities, then locked for held-out test evaluation. ECE was computed using 10 equal-width probability bins over [0, 1]. Calibration was applied in the 10-shot calibration condition without updating the underlying ranking model. Fine-tuning used the 10-shot support trials to adapt the model. Selective-prediction and coverage-risk calculations were configured in the codebase but are not central to the main manuscript claims.

### Reduced multiseed design

A reduced multiseed pass was performed for the decisive source conditions—pooled raw, pooled dataset-balanced, EEGMMIDB-only, and MILimbEEG-only—under zero-shot and 10-shot fine-tuning. Classical pipelines were effectively deterministic under frozen folds and support sets; EEGNet was repeated across multiple seeds and summarised in Table 4.

### Support-set resampling and threshold sensitivity

To assess whether the 10-shot conclusions depended on which support trials were selected, we performed 20 support-set draws per participant for the decisive classical conditions and summarised both mean metric values and per-participant support-draw standard deviations in Table 4.

Threshold sensitivity was evaluated by shifting the operating threshold by *±*0.05 and *±*0.10 around the locked value and recomputing balanced accuracy, sensitivity, specificity, ECE, Brier score, and F1; the resulting summaries are reported in Table 4.

### Sample-size-matched analysis

Because dataset balancing alters source size as well as source composition, we performed matched-source robustness comparisons between pooled raw and pooled dataset-balanced zero-shot transport. These comparisons are summarised in Tables 3 and 4.

### Sanity checks

Two sanity checks were prespecified.

- **Label permutation:** source labels were permuted and the zero-shot pipeline rerun.
- **Leakage rerun:** the pooled raw zero-shot pipeline was rerun under explicit leakage-control settings to confirm reproducibility of the leading result.

These analyses are summarised in Table 4.

### Metrics and statistical interpretation

Primary performance was AUROC. Secondary metrics were AUPRC, balanced accuracy, F1, Brier score, expected calibration error (ECE), sensitivity, and specificity. Because the translational question concerned participant-level transport, uncertainty was quantified at the participant level using bootstrap confidence intervals and gain intervals summarised in Table 4.

The study was designed for estimation and translational decision-making rather than formal winner-takes-all hypothesis testing. Accordingly, the main analyses emphasise point estimates, confidence intervals, gain intervals, and consistency across robustness analyses. Where conditions were compared without a formal equivalence or superiority test, the manuscript uses restrained language such as "numerically close" or "broadly tied".

### Software and reproducibility

Analyses were executed with locked YAML configurations for each experiment family, deterministic source-sampling controls, and scripted figure generation using the public lower-limb EEG transport repository. The analysis code, configuration files, and figure-generation scripts are publicly available in the referenced repository.[17]

### Use of large language models

A large language model was used during manuscript preparation for language editing and clarity-oriented restructuring only. It was not used for data analysis, code execution, numerical result generation, or statistical computation. All final text, interpretations, and submission decisions were reviewed and approved by the authors, who take responsibility for the manuscript.

## Results

### Task harmonisation and benchmark framing

We harmonised three public datasets to a common binary control variable, `effort_rest`, using fixed 2.0 s windows resampled to 125 Hz (Table 1). In EEGM- MIDB, both-feet execution and imagery events (`T2`) were mapped to effort and rest events (`T0`) to rest. In MILimbEEG, filename class 1 served as rest and lower-limb classes 4–7 were mapped to effort across execution and imagery modes, while intermediate non-target classes were excluded. In Stroke2025, event identifiers 3/4/5/6 were mapped to effort and 7/8 to a rest-proxy class in the locked benchmark. The common task was deliberately coarse: the study asked whether any defensible lower-limb effort-versus-rest signal transports, not whether finer subclass distinctions are yet viable.

The healthy source pool was internally distorted before transport. EEGMMIDB contributed a balanced source subset of 908 trials (454 rest and 454 effort), whereas MILimbEEG contributed 2520 trials with a markedly skewed class distribution (120 rest and 2400 effort). This mattered because the benchmark was designed not only to compare models, but also to test whether transport depends on how the healthy source is constructed. We therefore prespecified six source conditions: EEGMMIDB- only, MILimbEEG-only, pooled raw, pooled class-balanced, pooled dataset-balanced, and pooled class-plus-dataset-balanced.

The target transport analyses were aggregated over 27 participants with stroke, and the adaptation ladder was explicit: zero-shot transport, 10-shot calibration, and 10-shot fine-tuning, where 10-shot denotes 10 labelled trials per class. In the few-shot branch, support and test sets were trial-disjoint within participant but could be drawn from the same or different public Stroke2025 group labels; session/timepoint isolation was not enforced. The contribution was therefore not merely to ask whether healthy-to-stroke transfer exists; it was to provide a fully public lower-limb external benchmark n which source construction, minimal adaptation burden, and confound-sensitive montage controls were audited together under locked settings. Beyond the main ladder, the strengthening pass added participant-level bootstrap confidence intervals, reduced multiseed evaluation, 20-draw support-set resampling, sample-size-matched source comparisons, label-permutation checks, leakage reruns, and a within-dataset montage audit.

### Within-dataset benchmarking identifies a candidate regime but not a portable solution

Within-dataset performance on EEGMMIDB identified a candidate operating regime, but not a portable solution. Figure 5A and Table 5 summarise the within-dataset montage audit, which included the primary transport models plus an exploratory EEG Transformer comparator. In leave-one-subject-out EEGMMIDB analyses, EEGNet on the canonical 16-channel montage achieved the highest AUROC in the final bundle (0.782), with AUPRC 0.763 and balanced accuracy 0.714. On its own, that result could be read as encouraging.

However, the same montage audit already contained a warning. Frontal-control, motor-Laplacian, and temporal-control montages remained comparatively strong, with AUROCs of 0.735, 0.737, and 0.711, respectively. Within-dataset performance therefore identified a feasible regime for further translational testing, but it did not justify a portable or cleanly motor-specific claim. That distinction proved important once the analysis moved into healthy-to-stroke transport.

### Healthy-to-stroke zero-shot transport is weak

Within this benchmark, zero-shot healthy-to-stroke transport was weak across all model families. Figure 1A shows the main transport ladder, and Table 2 provides the numeric summary. In that ladder, CSP+LDA achieved the best zero-shot AUROC at 0.603, with AUPRC 0.579 and balanced accuracy 0.551. Riemann+TSLR trailed at AUROC 0.566. EEGNet remained near chance in the transport setting, with AUROC 0.527 and balanced accuracy 0.500.

**Fig. 1.**
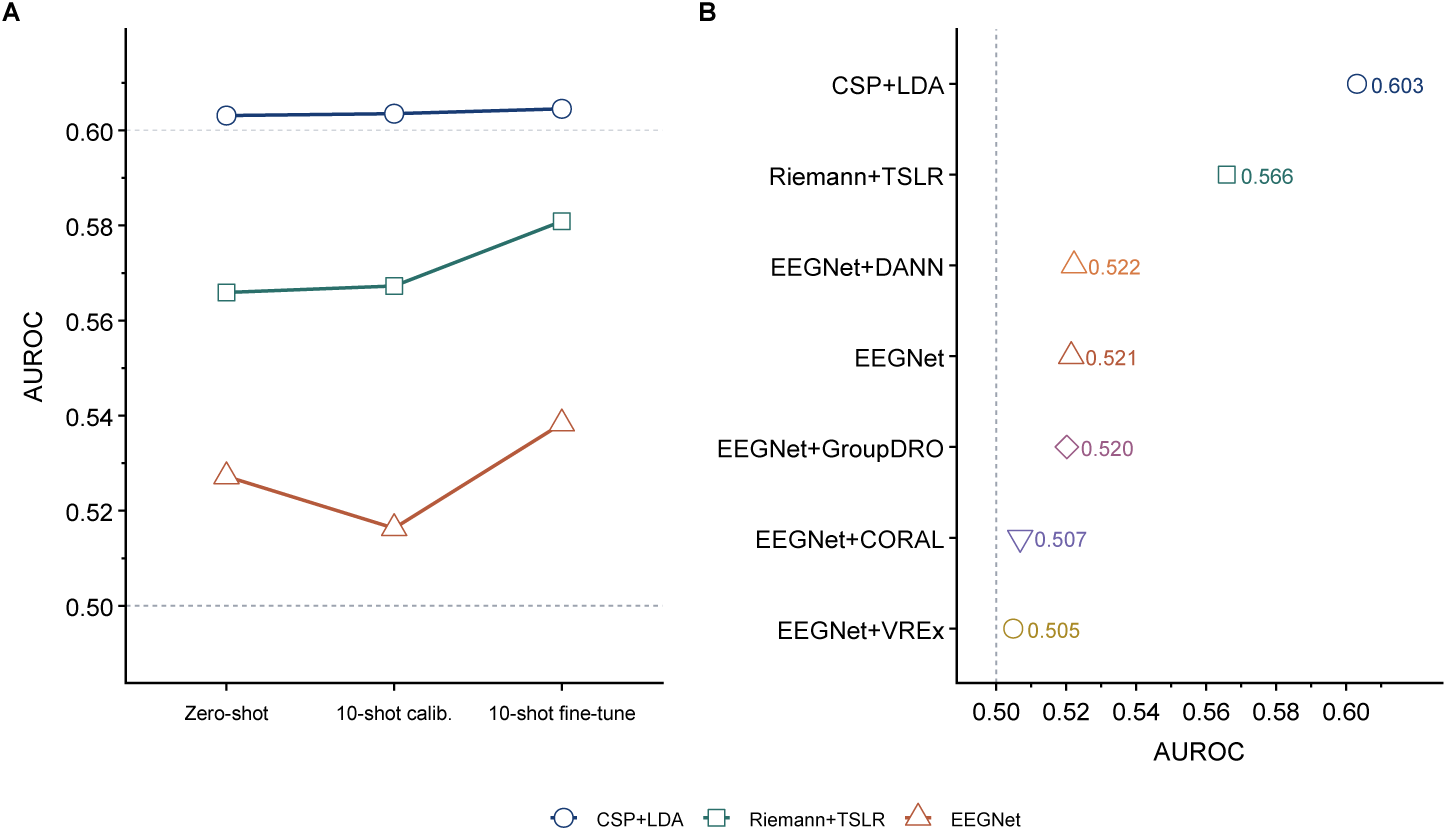
Healthy-to-stroke transport ladder. Panel A shows zero-shot, 10-shot calibration, and 10-shot fine-tune AUROC for CSP+LDA, Riemann+TSLR, and EEGNet. Panel B shows the domain-generalisation branch for the same transport benchmark. Classical baselines remained ahead of EEGNet, and adaptation primarily changed operating behaviour more than discrimination.

**Table 2.** Healthy-to-stroke transport ladder summary.

| Stage | Model | AU-ROC | AUPRC | Bal. acc. | Sens. | Spec. | ECE | Brier |
| --- | --- | --- | --- | --- | --- | --- | --- | --- |
| Zero-shot | CSP+LDA | 0.603 | 0.579 | 0.551 | 0.922 | 0.180 | 0.267 | 0.314 |
| Zero-shot | Riemann+TSLR | 0.566 | 0.553 | 0.547 | 0.815 | 0.279 | 0.209 | 0.298 |
| Zero-shot | EEGNet | 0.527 | 0.517 | 0.500 | 0.018 | 0.983 | 0.151 | 0.278 |
| 10-shot calibration | CSP+LDA | 0.604 | 0.580 | 0.568 | 0.652 | 0.485 | 0.035 | 0.249 |
| 10-shot calibration | Riemann+TSLR | 0.567 | 0.554 | 0.538 | 0.629 | 0.448 | 0.053 | 0.248 |
| 10-shot calibration | EEGNet | 0.516 | 0.512 | 0.513 | 0.578 | 0.448 | 0.051 | 0.253 |
| 10-shot fine-tune | CSP+LDA | 0.605 | 0.581 | 0.566 | 0.630 | 0.503 | 0.045 | 0.249 |
| 10-shot fine-tune | Riemann+TSLR | 0.581 | 0.568 | 0.553 | 0.585 | 0.522 | 0.073 | 0.251 |
| 10-shot fine-tune | EEGNet | 0.538 | 0.527 | 0.522 | 0.647 | 0.397 | 0.036 | 0.251 |
| Domain generalization | CSP+LDA | 0.603 | 0.579 | 0.551 | 0.922 | 0.180 | 0.267 | 0.314 |
| Domain generalization | Riemann+TSLR | 0.566 | 0.553 | 0.547 | 0.815 | 0.279 | 0.209 | 0.298 |
| Domain generalization | EEGNet | 0.521 | 0.515 | 0.500 | 0.034 | 0.966 | 0.146 | 0.276 |
| Domain generalization | EEGNet+DANN | 0.522 | 0.518 | 0.501 | 0.047 | 0.956 | 0.157 | 0.280 |
| Domain generalization | EEGNet+GroupDRO | 0.520 | 0.514 | 0.495 | 0.045 | 0.944 | 0.150 | 0.280 |
| Domain generalization | EEGNet+CORAL | 0.507 | 0.507 | 0.499 | 0.038 | 0.960 | 0.169 | 0.286 |
| Domain generalization | EEGNet+VREx | 0.505 | 0.509 | 0.499 | 0.063 | 0.935 | 0.157 | 0.285 |

Note: Main ladder values are point estimates from the locked transport analysis. To foreground uncertainty, the matched-source robustness pass yielded zero-shot CSP+LDA AUROC 0.609 (95% CI 0.584–0.635) for pooled raw and 0.603 (95% CI 0.579–0.628) for pooled dataset-balanced, with corresponding zero-shot to 10-shot fine-tune delta AUROC 0.001 (95% CI -0.001 to 0.003) and 0.001 (95% CI -0.000 to 0.003), respectively (Table 4). AUROC, area under the receiver operating characteristic curve; AUPRC, area under the precision-recall curve; Bal. acc., balanced accuracy; ECE, expected calibration error.

The operating behaviour of the leading classical and deep models diverged sharply. CSP+LDA in zero-shot mode was heavily sensitivity-weighted (sensitivity 0.922; specificity 0.180), whereas EEGNet was almost entirely negative-predicting at the chosen operating point (sensitivity 0.018; specificity 0.983). For CSP+LDA, the weakness was partly an operating-point and calibration problem. For EEGNet, the problem extended to rank-order discrimination itself, because AUROC remained low even before threshold choice was considered.

Bootstrap confidence intervals from the matched-source robustness pass were consistent with this interpretation. In the sample-size-matched pooled raw comparison, zero-shot CSP+LDA yielded AUROC 0.609 (95% CI 0.584–0.635); the matched pooled dataset-balanced comparison yielded AUROC 0.603 (0.579–0.628) (Table 4). Neither condition approached a clinically reassuring transport regime.

### Calibration and 10-shot adaptation improve operating behaviour more than discrimination

Minimal adaptation helped, but it helped mainly by changing operating behaviour rather than improving rank-order discrimination. Figure 1A and Figure 3B show this separation between discrimination and operating behaviour, with the underlying values summarised in Table 2. In the main transport ladder, 10-shot temperature calibration left CSP+LDA AUROC essentially unchanged (0.603 to 0.604) while reducing ECE from 0.267 to 0.035 and increasing specificity from 0.180 to 0.485. Sensitivity fell from 0.922 to 0.652 as the operating point moved away from an almost-all-positive classifier. The principal effect of calibration was therefore to change how the model behaved, not what it fundamentally ranked.

Ten-shot fine-tuning produced only modest AUROC gains. Figure 3A visualises those gain estimates across source conditions, while Table 2 and Table 4 provide the stage-wise and bootstrap summaries. In the main transport ladder, CSP+LDA improved from 0.603 to 0.605, Riemann+TSLR from 0.566 to 0.581, and EEGNet from 0.527 to 0.538. Bootstrap gain estimates reinforced the modesty of those changes. For pooled raw CSP+LDA, the mean AUROC gain from zero-shot to 10-shot fine-tuning was 0.001 (95% CI −0.001 to 0.003); for pooled dataset-balanced CSP+LDA it was 0.001 (−0.000 to 0.003). Riemann+TSLR gained more than CSP+LDA under the same conditions, but its absolute performance remained below the leading classical result. Operating point remained fragile to threshold choice even when AUROC did not move. In the sample-size-matched pooled dataset-balanced zero-shot CSP+LDA analysis, balanced accuracy ranged from 0.499 to 0.562 as the threshold was shifted by *±*0.10. Over the same range, sensitivity varied from 0.010 to 0.983 and specificity from 0.036 to 0.990, while AUROC remained fixed at 0.603 (Table 4). The translational implication is straightforward: discrimination, calibration, and threshold policy are separate variables, and bedside viability depends as much on the latter two as on AUROC.

### Source construction matters, but no source recipe fully rescues transport

Source construction shaped transport, but no retrospective source recipe rescued it. Figure 2 summarises the source-ablation matrix, and Table 3 provides the numeric values. MILimbEEG-only was the clearest failure condition. In zero-shot transport, MILimbEEG-only CSP+LDA reached AUROC 0.463, with balanced accuracy 0.499, sensitivity 0.006, and specificity 0.992. The corresponding 10-shot fine-tuned result remained poor at AUROC 0.468. This pattern is difficult to explain as simple seed variance. It points to source mismatch, probably compounded by the source pool’s extreme class imbalance and task semantics.

**Fig. 2.**
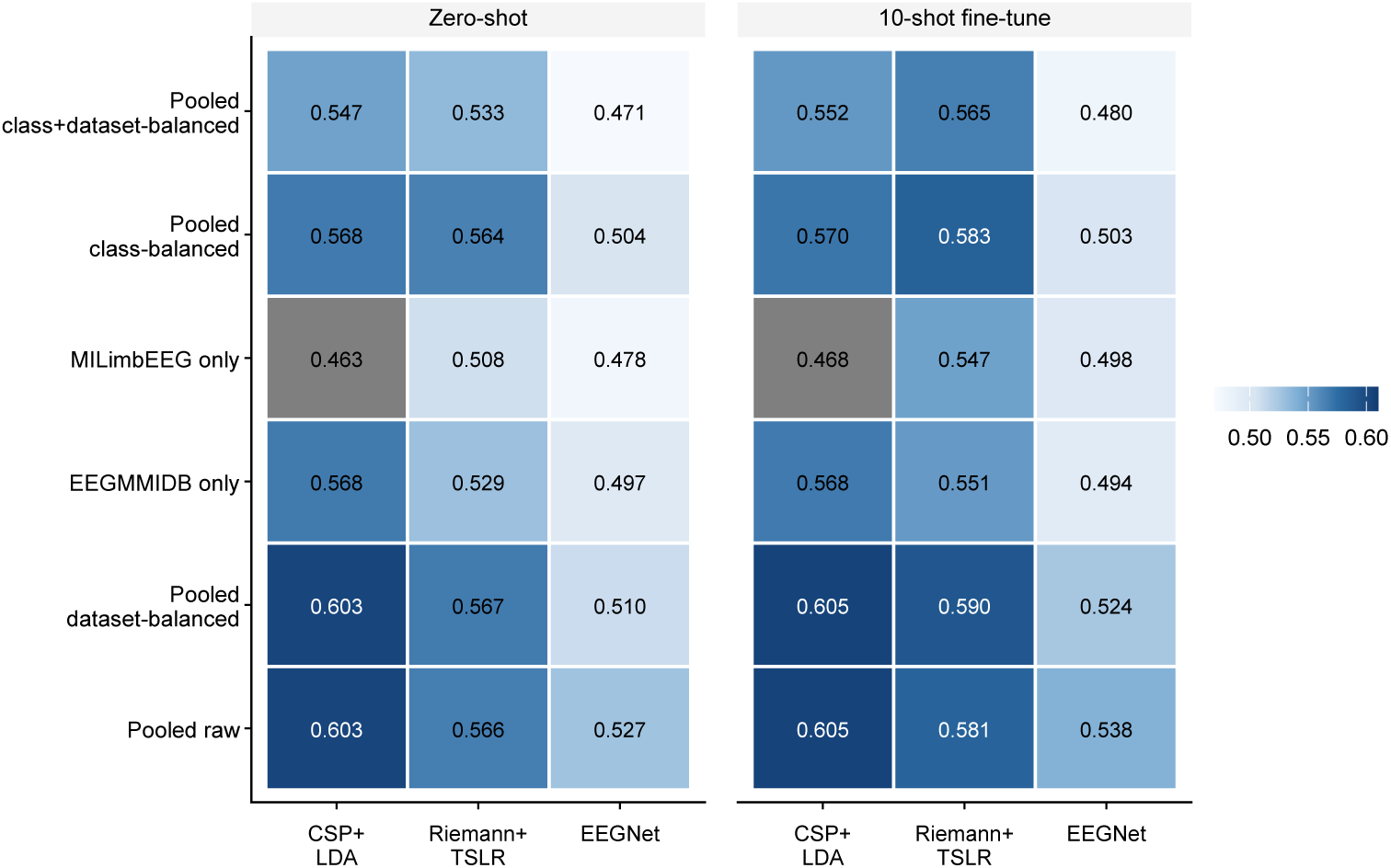
Source-ablation AUROC heatmap. Left, zero-shot; right, 10-shot fine-tune. AUROC is shown across six source-construction conditions: EEGMMIDB-only, MILimbEEG-only, pooled raw, pooled class-balanced, pooled dataset-balanced, and pooled class-plus-dataset-balanced. MILimbEEG-only was consistently weak, whereas pooled raw and pooled dataset-balanced sources remained broadly tied for the classical lead.

**Fig. 3.**
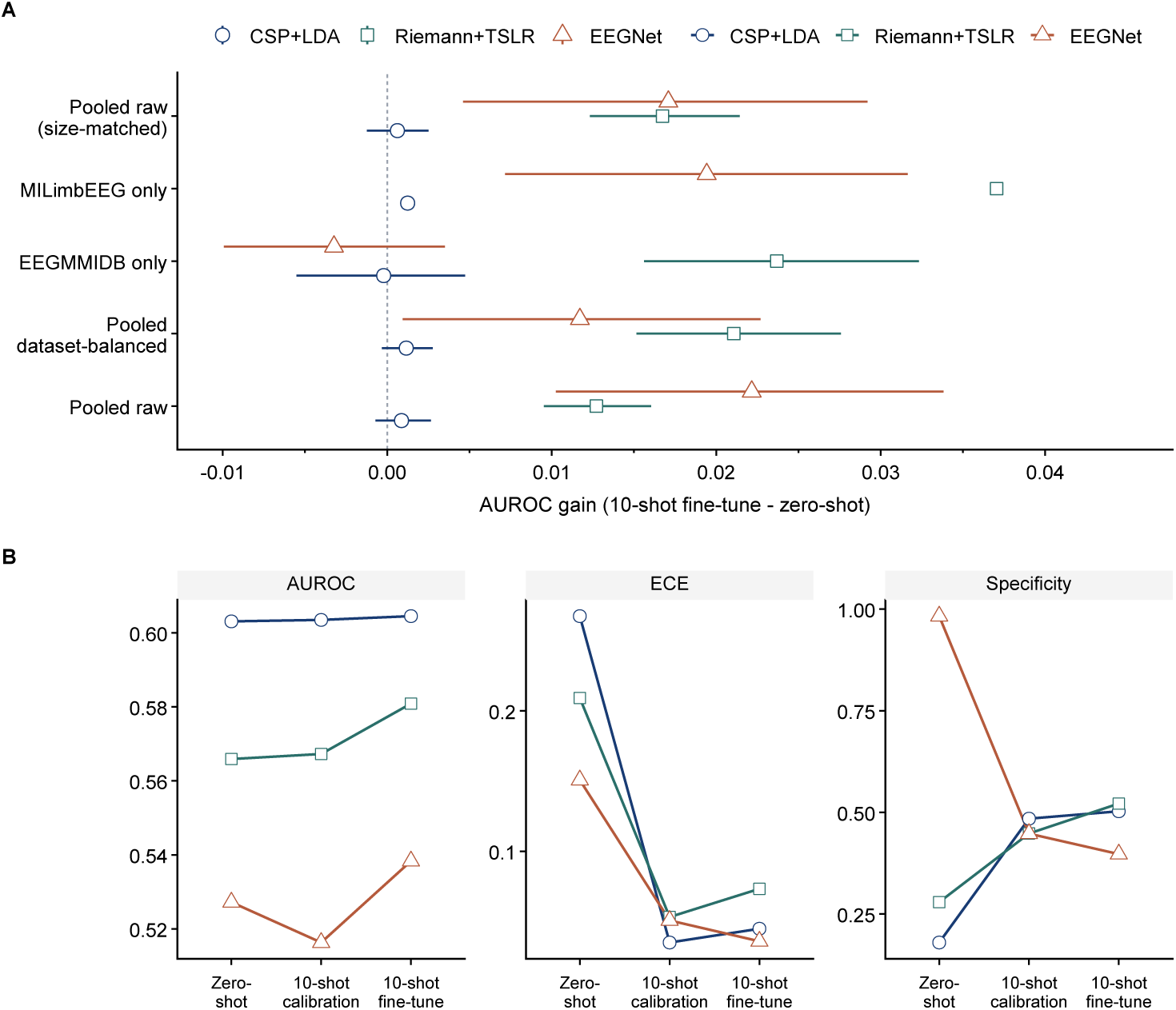
AUROC gain and operating-behaviour profiles. Panel A shows bootstrap mean AUROC gain with 95% confidence intervals from zero-shot to 10-shot fine-tuning across source conditions and models. Panel B shows AUROC, expected calibration error, and specificity across zero-shot, calibration, and fine-tune stages. Minimal adaptation yielded only modest discriminative gain and mainly shifted operating behaviour.

**Table 3:**
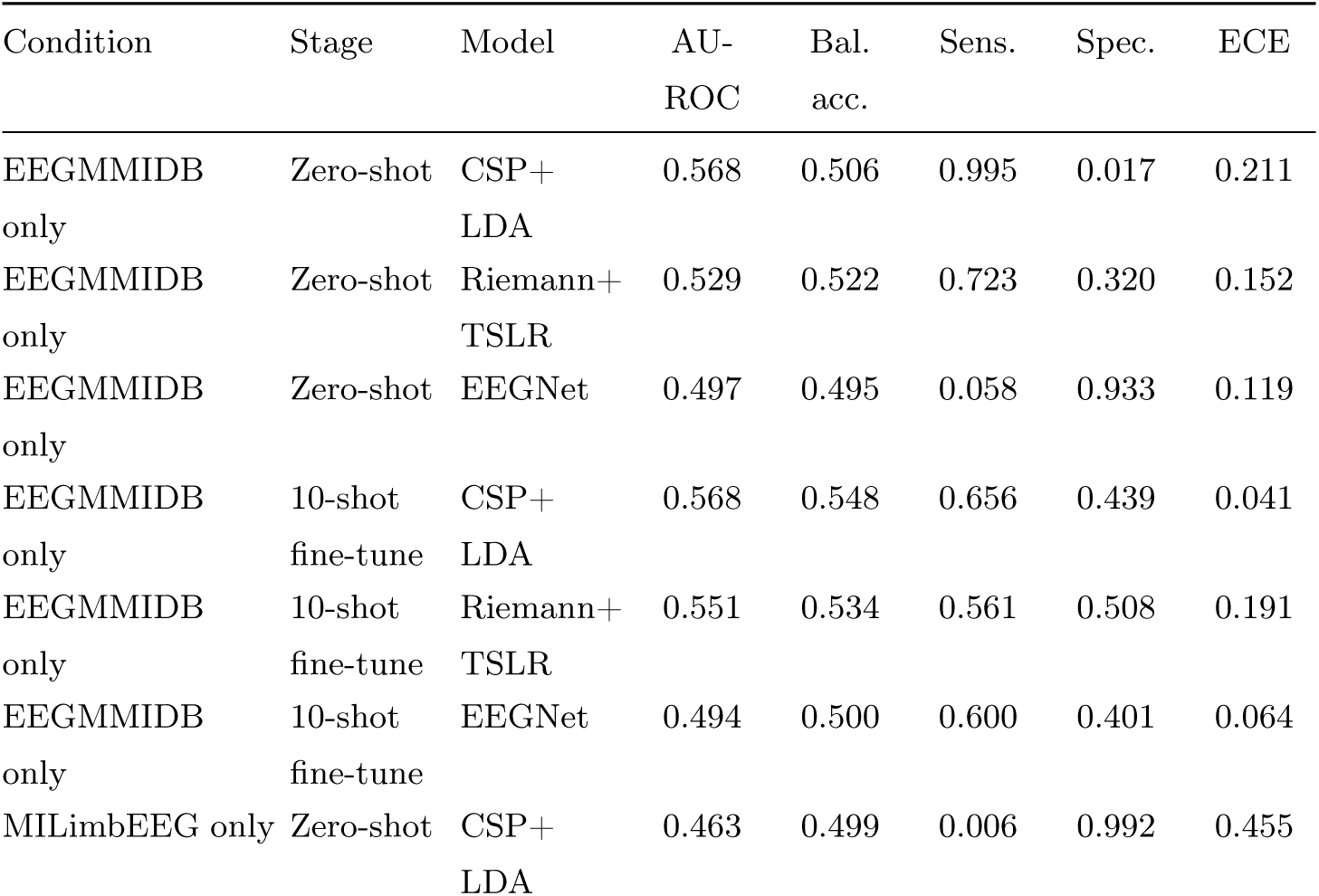
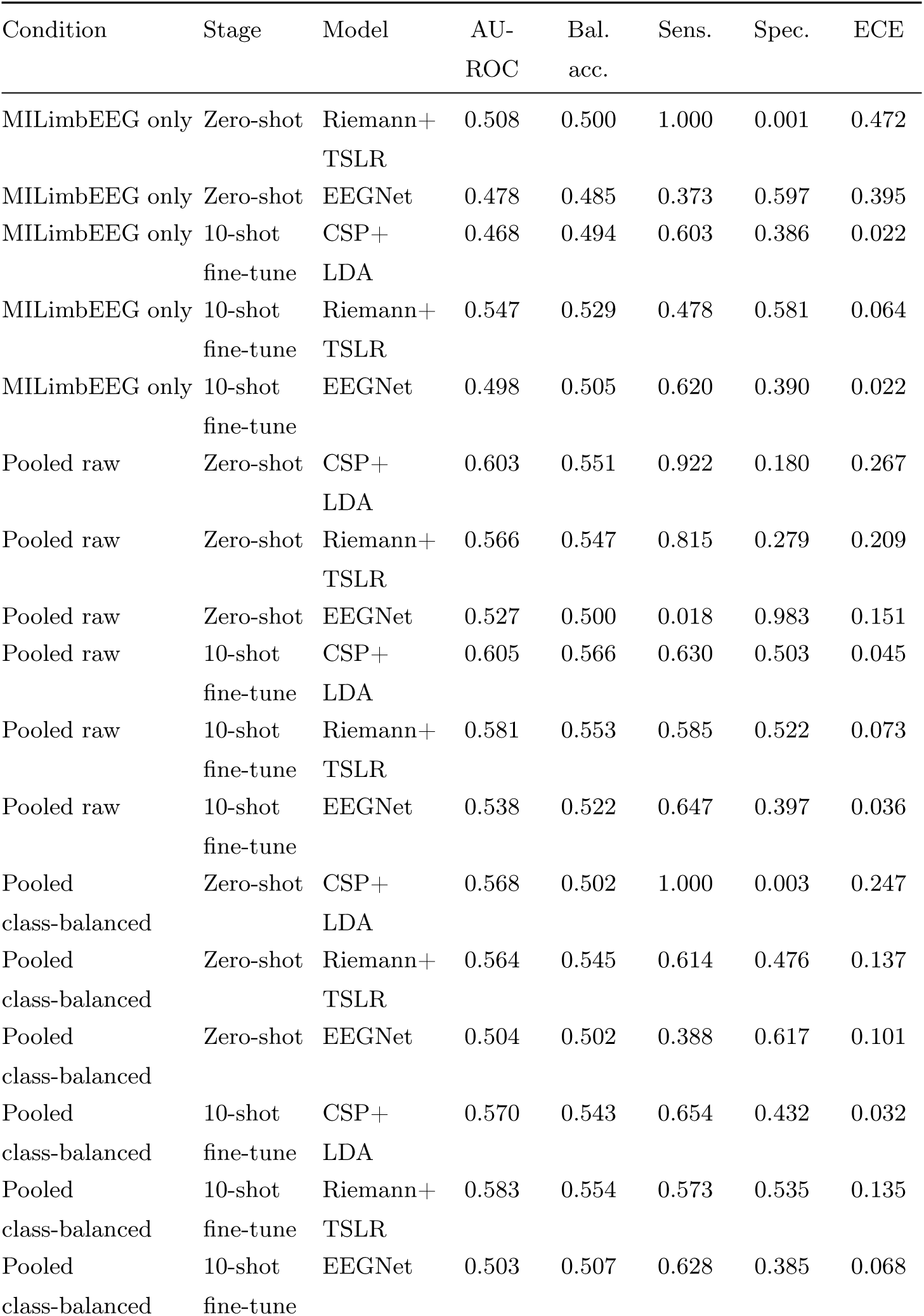
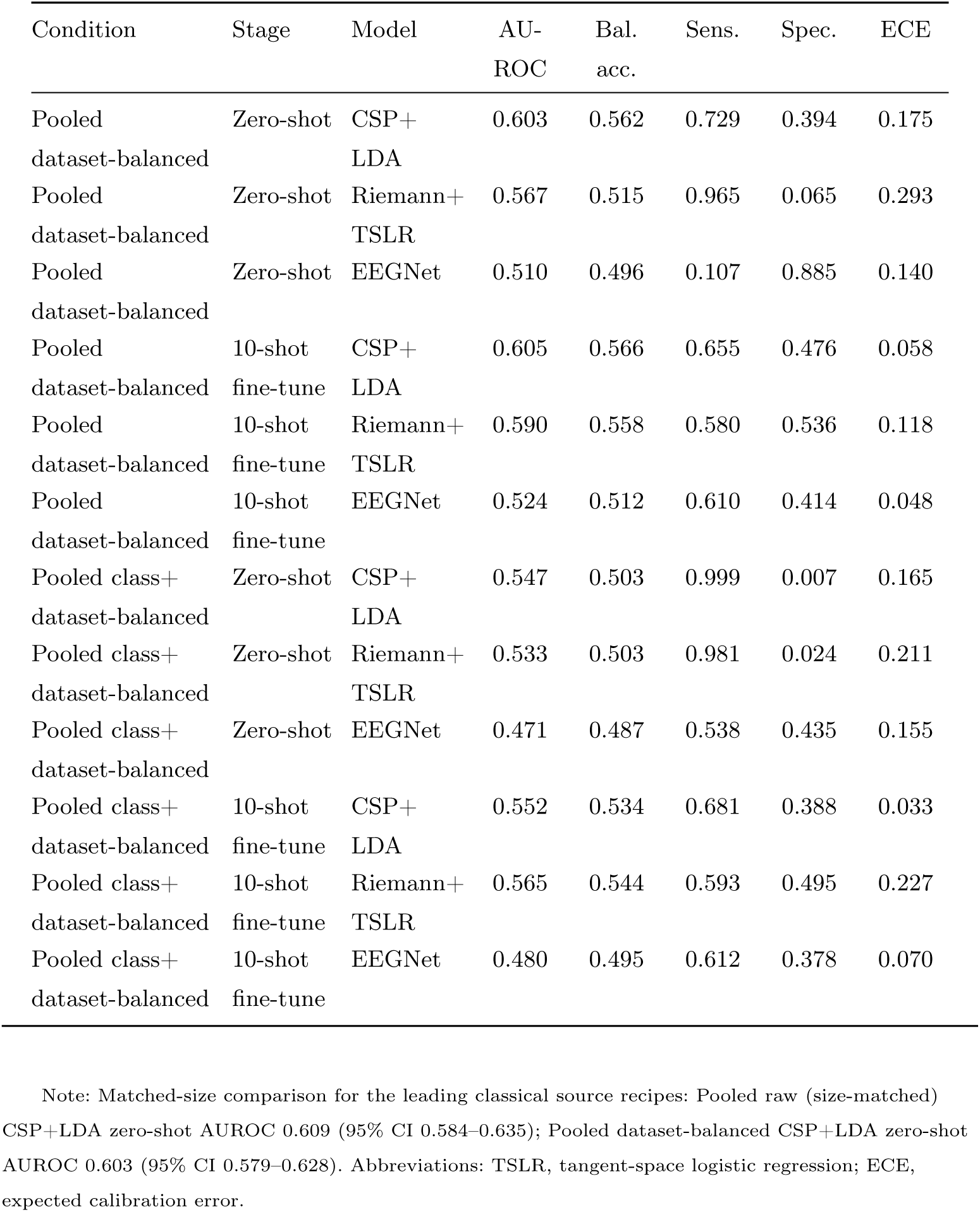
Source-ablation and matched-source analyses.

| Condition | Stage | Model | AU-ROC | Bal. acc. | Sens. | Spec. | ECE |
| --- | --- | --- | --- | --- | --- | --- | --- |
| EEGMMIDB only | Zero-shot | CSP+<br>LDA | 0.568 | 0.506 | 0.995 | 0.017 | 0.211 |
| EEGMMIDB only | Zero-shot | Riemann+<br>TSLR | 0.529 | 0.522 | 0.723 | 0.320 | 0.152 |
| EEGMMIDB only | Zero-shot | EEGNet | 0.497 | 0.495 | 0.058 | 0.933 | 0.119 |
| EEGMMIDB only | 10-shot<br>fine-tune | CSP+<br>LDA | 0.568 | 0.548 | 0.656 | 0.439 | 0.041 |
| EEGMMIDB only | 10-shot<br>fine-tune | Riemann+<br>TSLR | 0.551 | 0.534 | 0.561 | 0.508 | 0.191 |
| EEGMMIDB only | 10-shot<br>fine-tune | EEGNet | 0.494 | 0.500 | 0.600 | 0.401 | 0.064 |
| MILimbEEG only | Zero-shot | CSP+<br>LDA | 0.463 | 0.499 | 0.006 | 0.992 | 0.455 |
| MILimbEEG only | Zero-shot | Riemann+ | 0.508 | 0.500 | 1.000 | 0.001 | 0.472 |
|  |  | TSLR |  |  |  |  |  |
| MILimbEEG only | Zero-shot | EEGNet | 0.478 | 0.485 | 0.373 | 0.597 | 0.395 |
| MILimbEEG only | 10-shot | CSP+ | 0.468 | 0.494 | 0.603 | 0.386 | 0.022 |
|  | fine-tune | LDA |  |  |  |  |  |
| MILimbEEG only | 10-shot | Riemann+ | 0.547 | 0.529 | 0.478 | 0.581 | 0.064 |
|  | fine-tune | TSLR |  |  |  |  |  |
| MILimbEEG only | 10-shot | EEGNet | 0.498 | 0.505 | 0.620 | 0.390 | 0.022 |
|  | fine-tune |  |  |  |  |  |  |
| Pooled raw | Zero-shot | CSP+ | 0.603 | 0.551 | 0.922 | 0.180 | 0.267 |
|  |  | LDA |  |  |  |  |  |
| Pooled raw | Zero-shot | Riemann+ | 0.566 | 0.547 | 0.815 | 0.279 | 0.209 |
|  |  | TSLR |  |  |  |  |  |
| Pooled raw | Zero-shot | EEGNet | 0.527 | 0.500 | 0.018 | 0.983 | 0.151 |
| Pooled raw | 10-shot | CSP+ | 0.605 | 0.566 | 0.630 | 0.503 | 0.045 |
|  | fine-tune | LDA |  |  |  |  |  |
| Pooled raw | 10-shot | Riemann+ | 0.581 | 0.553 | 0.585 | 0.522 | 0.073 |
|  | fine-tune | TSLR |  |  |  |  |  |
| Pooled raw | 10-shot | EEGNet | 0.538 | 0.522 | 0.647 | 0.397 | 0.036 |
|  | fine-tune |  |  |  |  |  |  |
| Pooled | Zero-shot | CSP+ | 0.568 | 0.502 | 1.000 | 0.003 | 0.247 |
| class-balanced |  | LDA |  |  |  |  |  |
| Pooled | Zero-shot | Riemann+ | 0.564 | 0.545 | 0.614 | 0.476 | 0.137 |
| class-balanced |  | TSLR |  |  |  |  |  |
| Pooled | Zero-shot | EEGNet | 0.504 | 0.502 | 0.388 | 0.617 | 0.101 |
| class-balanced |  |  |  |  |  |  |  |
| Pooled | 10-shot | CSP+ | 0.570 | 0.543 | 0.654 | 0.432 | 0.032 |
| class-balanced | fine-tune | LDA |  |  |  |  |  |
| Pooled | 10-shot | Riemann+ | 0.583 | 0.554 | 0.573 | 0.535 | 0.135 |
| class-balanced | fine-tune | TSLR |  |  |  |  |  |
| Pooled | 10-shot | EEGNet | 0.503 | 0.507 | 0.628 | 0.385 | 0.068 |
| class-balanced | fine-tune |  |  |  |  |  |  |
| Pooled | Zero-shot | CSP+ | 0.603 | 0.562 | 0.729 | 0.394 | 0.175 |
| dataset-balanced |  | LDA |  |  |  |  |  |
| Pooled | Zero-shot | Riemann+ | 0.567 | 0.515 | 0.965 | 0.065 | 0.293 |
| dataset-balanced |  | TSLR |  |  |  |  |  |
| Pooled | Zero-shot | EEGNet | 0.510 | 0.496 | 0.107 | 0.885 | 0.140 |
| dataset-balanced |  |  |  |  |  |  |  |
| Pooled | 10-shot | CSP+ | 0.605 | 0.566 | 0.655 | 0.476 | 0.058 |
| dataset-balanced | fine-tune | LDA |  |  |  |  |  |
| Pooled | 10-shot | Riemann+ | 0.590 | 0.558 | 0.580 | 0.536 | 0.118 |
| dataset-balanced | fine-tune | TSLR |  |  |  |  |  |
| Pooled | 10-shot | EEGNet | 0.524 | 0.512 | 0.610 | 0.414 | 0.048 |
| dataset-balanced | fine-tune |  |  |  |  |  |  |
| Pooled class+ | Zero-shot | CSP+ | 0.547 | 0.503 | 0.999 | 0.007 | 0.165 |
| dataset-balanced |  | LDA |  |  |  |  |  |
| Pooled class+ | Zero-shot | Riemann+ | 0.533 | 0.503 | 0.981 | 0.024 | 0.211 |
| dataset-balanced |  | TSLR |  |  |  |  |  |
| Pooled class+ | Zero-shot | EEGNet | 0.471 | 0.487 | 0.538 | 0.435 | 0.155 |
| dataset-balanced |  |  |  |  |  |  |  |
| Pooled class+ | 10-shot | CSP+ | 0.552 | 0.534 | 0.681 | 0.388 | 0.033 |
| dataset-balanced | fine-tune | LDA |  |  |  |  |  |
| Pooled class+ | 10-shot | Riemann+ | 0.565 | 0.544 | 0.593 | 0.495 | 0.227 |
| dataset-balanced | fine-tune | TSLR |  |  |  |  |  |
| Pooled class+ | 10-shot | EEGNet | 0.480 | 0.495 | 0.612 | 0.378 | 0.070 |
| dataset-balanced | fine-tune |  |  |  |  |  |  |

Note: Matched-size comparison for the leading classical source recipes: Pooled raw (size-matched) CSP+LDA zero-shot AUROC 0.609 (95% CI 0.584–0.635); Pooled dataset-balanced CSP+LDA zero-shot AUROC 0.603 (95% CI 0.579–0.628). Abbreviations: TSLR, tangent-space logistic regression; ECE, expected calibration error.

By contrast, pooled raw and pooled dataset-balanced sources were broadly tied for the classical lead. In zero-shot mode, pooled raw CSP+LDA achieved AUROC 0.603 and pooled dataset-balanced CSP+LDA 0.603. After 10-shot fine-tuning, pooled raw CSP+LDA reached 0.6045 and pooled dataset-balanced CSP+LDA 0.6047. Supplementary Figure S2 shows the matched-source comparison, which did not materially change that conclusion, supporting a restrained reading: dataset balancing may help interpretability of source composition, but it did not materially alter discriminative transport in this retrospective benchmark.

Aggressive class-plus-dataset balancing reduced source size too far to help. The fully balanced pooled source shrank to 480 training trials (120 rest and 120 effort from each source dataset) and underperformed the broader pooled strategies, with CSP+LDA AUROC 0.547 in zero-shot mode and 0.552 after 10-shot fine-tuning (Table 3). This cautions against simplistic balancing assumptions: source construction matters, but transport is not rescued by indiscriminate subsampling.

Deep models did not provide the main performance gains in this source-ablation matrix. The best EEGNet transport result in the final matrix was pooled raw 10-shot fine-tuning at AUROC 0.538. Figure 1B shows that exploratory domain-generalised variants did not change that picture; the best deep DG branch, EEGNet+DANN, reached AUROC 0.522 in the main transport ladder, still well below the classical baselines in Table 2.

### Support-set resampling, bootstrap confidence intervals, and sanity checks reinforce the main conclusion

The strengthening analyses hardened the translational interpretation rather than changing it. Figure 4A-B summarises the reduced multiseed and support-resampling results, and Table 4 provides the corresponding numerical summaries. Twenty-draw support-set resampling showed that AUROC was relatively stable within each classical condition, but operating point was not. For pooled raw CSP+LDA, mean AUROC across support draws was 0.604, with mean per-subject support-draw AUROC standard deviation 0.005. Yet mean support-draw standard deviation was 0.235 for sensitivity and 0.220 for specificity. Pooled dataset-balanced CSP+LDA showed the same pattern: mean AUROC 0.604, AUROC support-draw standard deviation 0.005, but sensitivity and specificity standard deviations of 0.229 and 0.218, respectively. In translational terms, support-set choice affected how the model operated much more than how it ranked.

**Fig. 4.**
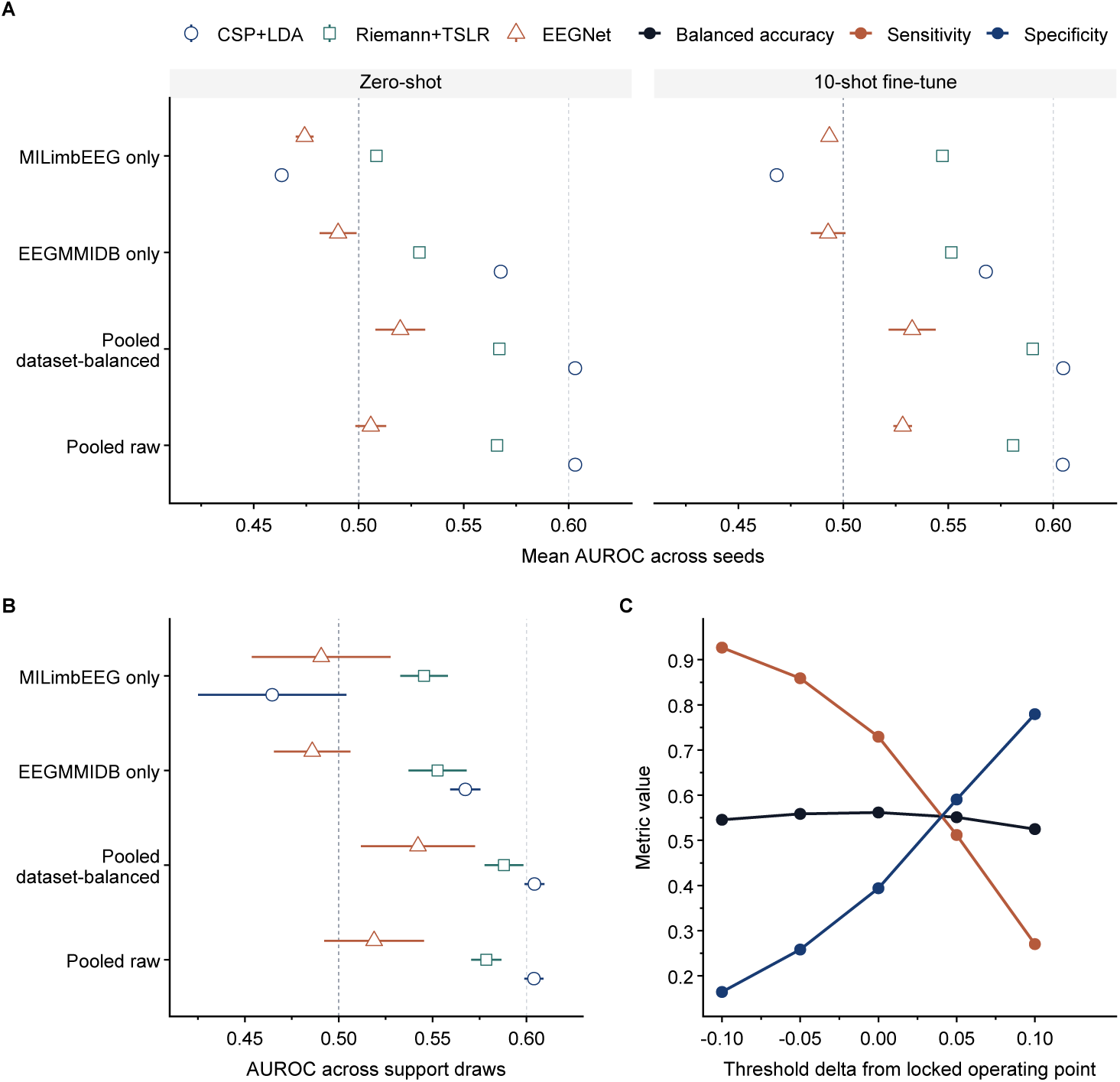
Robustness analyses. Panel A shows reduced multiseed mean AUROC across key source conditions. Panel B shows support-set resampling stability. Panel C shows threshold sensitivity for pooled dataset-balanced zero-shot CSP+LDA. AUROC was comparatively stable, whereas sensitivity and specificity remained much more dependent on support-set choice and threshold policy.

**Table 4.** Robustness and sanity checks.

| Analysis | Condition | Model | Key result | Interval /variability |
| --- | --- | --- | --- | --- |
| Bootstrap CI | Pooled raw (size-matched) | CSP+<br>LDA | AUROC<br>0.609 | 95% CI 0.584-0.635 |
| Bootstrap CI | Pooled raw (size-matched) | EEGNet | AUROC<br>0.509 | 95% CI 0.486-0.534 |
| Bootstrap CI | Pooled dataset-balanced | CSP+<br>LDA | AUROC<br>0.603 | 95% CI 0.579-0.628 |
| Bootstrap CI | Pooled dataset-balanced | EEGNet | AUROC<br>0.531 | 95% CI 0.511-0.551 |
| Label permutation | Label permutation control | CSP+<br>LDA | AUROC<br>0.488 | 95% CI 0.465-0.511 |
| Label permutation | Label permutation control | EEGNet | AUROC<br>0.493 | 95% CI 0.476-0.509 |
| Leakage rerun | Leakage rerun control | CSP+<br>LDA | AUROC<br>0.603 | 95% CI 0.579-0.629 |
| Leakage rerun | Leakage rerun control | EEGNet | AUROC<br>0.497 | 95% CI 0.476-0.518 |
| Bootstrap gain | Pooled raw | CSP+<br>LDA | Delta<br>AUROC<br>0.001 | 95% CI -0.001-0.003 |
| Bootstrap gain | Pooled dataset-balanced | CSP+<br>LDA | Delta<br>AUROC<br>0.001 | 95% CI -0.000-0.003 |
| Support-set resampling | Pooled raw (10-shot fine-tune) | CSP+<br>LDA | AUROC<br>0.604 | AUROC SDmean 0.005; Sens SDmean 0.235; Spec SDmean 0.220 |
| Support-set resampling | Pooled dataset-balanced (10-shot fine-tune) | CSP+<br>LDA | AUROC<br>0.604 | AUROC SDmean 0.005; Sens SDmean 0.229; Spec SDmean 0.218 |
| Threshold sensitivity | Pooled dataset-balanced (zero-shot) | CSP+<br>LDA | Bal. acc.<br>0.546 /<br>0.562 /<br>0.525 | Delta -0.10 /0 /+0.10; Sens 0.927→0.270; Spec 0.164→0.779 |
| Reduced multiseed | Pooled raw (zero-shot) | EEGNet | AUROC<br>0.506 ±<br>0.007 | Bal. acc. 0.498 ± 0.003 |
| Reduced multiseed | Pooled dataset-balanced (zero-shot) | EEGNet | AUROC<br>0.520 ±<br>0.012 | Bal. acc. 0.498 ± 0.006 |

Note: CI, confidence interval; Bal. acc., balanced accuracy; SDmean, mean within-participant standard deviation across support draws.

Reduced multiseed analysis did not change the qualitative story. Classical models were effectively deterministic under frozen target folds and support splits. EEGNet varied modestly but remained weak: pooled dataset-balanced zero-shot AUROC was 0.520 *±* 0.012 and pooled dataset-balanced 10-shot fine-tuned AUROC 0.533 *±* 0.011; pooled raw zero-shot AUROC was 0.506 *±* 0.007 and pooled raw 10-shot fine-tuned AUROC 0.528 *±* 0.004 (Table 4). The conclusion that source construction mattered more than compact deep-model novelty therefore remained stable under seed perturbation.

Sanity checks supported that the negative result was real. Supplementary Figure S1 and Table 4 summarise those controls. Under label permutation, performance fell to chance-like values: CSP+LDA AUROC 0.488 (95% CI 0.465–0.511) and EEGNet 0.493 (0.476–0.509). A leakage rerun reproduced the leading classical result in the pooled raw zero-shot setting, with CSP+LDA AUROC 0.603 (0.579–0.629). The main transport failure was therefore not plausibly explained by leakage or labelling artefact. Additional file 1 contains Supplementary Figures S1 and S2, which provide the label-permutation and leakage-control checks and the matched-source pooled raw versus pooled dataset-balanced comparison.

### Physiology audit limits a motor-specific interpretation

The physiology audit placed an explicit limit on interpretation. Figure 5A-B and Table 5 summarise these montage-control analyses. In within-dataset EEGMMIDB analyses, EEGNet on the canonical 16-channel montage achieved AUROC 0.782. However, frontal-control and motor-Laplacian montages were both around 0.735–0.737, and temporal-control remained 0.711. This does not show that the signal is artifactual or unusable. It does show that the present retrospective evidence does not support a strong claim that transport-relevant information arose predominantly from a narrowly motor-cortical source.

**Fig. 5.**
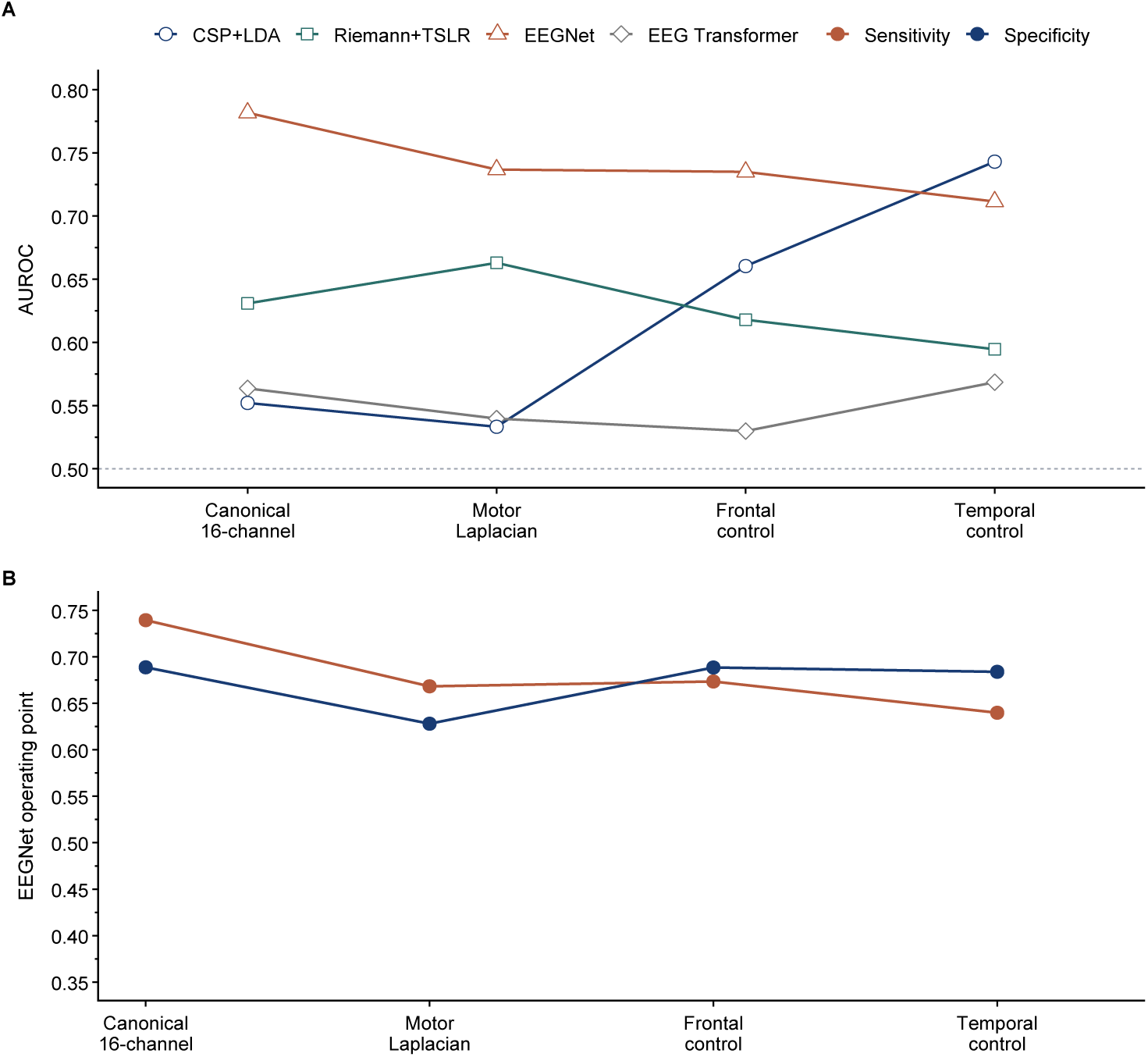
Physiology audit across montage controls. Panel A shows leave-one-subject-out AUROC across canonical, motor-Laplacian, frontal-control, and temporal-control montages. Panel B shows EEGNet sensitivity and specificity across the same montage controls. Competitive frontal and temporal control performance limits a strong motor-cortex-specific interpretation of the current retrospective signal.

**Table 5.** Physiology and montage audit.

| Montage | Model | AU-ROC | AUPRC | Bal. acc. | Sens. | Spec. | ECE |
| --- | --- | --- | --- | --- | --- | --- | --- |
| Canonical | CSP+LDA | 0.552 | 0.548 | 0.485 | 0.539 | 0.431 | 0.285 |
| 16-channel |  |  |  |  |  |  |  |
| Canonical | Riemann+TSLR | 0.631 | 0.634 | 0.518 | 0.521 | 0.514 | 0.327 |
| 16-channel |  |  |  |  |  |  |  |
| Canonical | EEGNet | 0.782 | 0.763 | 0.714 | 0.740 | 0.689 | 0.177 |
| 16-channel |  |  |  |  |  |  |  |
| Canonical | EEG | 0.564 | 0.575 | 0.541 | 0.615 | 0.467 | 0.145 |
| 16-channel | Transformer |  |  |  |  |  |  |
| Motor Laplacian | CSP+LDA | 0.533 | 0.549 | 0.485 | 0.512 | 0.457 | 0.212 |
| Motor Laplacian | Riemann+TSLR | 0.663 | 0.670 | 0.588 | 0.461 | 0.714 | 0.176 |
| Motor Laplacian | EEGNet | 0.737 | 0.745 | 0.648 | 0.668 | 0.628 | 0.179 |
| Motor Laplacian | EEG | 0.540 | 0.554 | 0.524 | 0.629 | 0.418 | 0.171 |
|  | Transformer |  |  |  |  |  |  |
| Frontal control | CSP+LDA | 0.660 | 0.651 | 0.607 | 0.541 | 0.673 | 0.212 |
| Frontal control | Riemann+TSLR | 0.618 | 0.617 | 0.603 | 0.581 | 0.625 | 0.237 |
| Frontal control | EEGNet | 0.735 | 0.739 | 0.681 | 0.674 | 0.689 | 0.139 |
| Frontal control | EEG | 0.530 | 0.556 | 0.513 | 0.548 | 0.478 | 0.148 |
|  | Transformer |  |  |  |  |  |  |
| Temporal control | CSP+LDA | 0.743 | 0.724 | 0.552 | 0.704 | 0.400 | 0.300 |
| Temporal control | Riemann+TSLR | 0.595 | 0.600 | 0.500 | 0.523 | 0.476 | 0.348 |
| Temporal control | EEGNet | 0.711 | 0.709 | 0.662 | 0.640 | 0.684 | 0.175 |
| Temporal control | EEG | 0.568 | 0.582 | 0.515 | 0.387 | 0.643 | 0.195 |
|  | Transformer |  |  |  |  |  |  |

Note: AUPRC, area under the precision-recall curve; Bal. acc., balanced accuracy; ECE, expected calibration error.

That limitation matters for translation. If frontal and temporal controls remain relatively competitive in retrospective public data, then claims about brain-specificity, validated digital biomarkers, or readiness for restorative neurotechnology should be made cautiously. The correct implication is methodological: future validation needs harmonised acquisition with explicit EOG, EMG, and kinematic confound channels, not stronger rhetoric about cortical specificity.

## Discussion

This study addressed a translational question that within-dataset EEG benchmarks cannot answer: whether lower-limb effort-versus-rest decoders learned in healthy public datasets survive transport into stroke. Within this public benchmark, transport remained weak. Across three public datasets harmonised to a common control variable, healthy-to-stroke zero-shot transport was weak, minimal adaptation yielded only modest gains in discrimination, and compact deep models did not outperform classical baselines. The contribution is not the abstract idea of healthy-to-stroke transfer alone, but a public external benchmark in which source construction, adaptation burden, and confound-sensitive montage controls were audited together under locked conditions. The principal message is therefore not that lower-limb EEG decoding is impossible, but that, within this benchmark, the main bottlenecks lay in source construction, adaptation burden, calibration, and unresolved confound sensitivity.

That is a clinically relevant result. Lower-limb EEG control continues to attract interest because it may eventually contribute to neurorehabilitation, closed-loop feedback, and non-invasive brain–spine interface strategies.[1–4] Yet those future applications require a signal that survives the move from healthy training data to stroke-domain use. A paper that directly examines that move—and shows that it is currently weak—is more useful to translational neuroengineering than another healthy within-dataset leaderboard. Careful external evaluations are part of how translational neuroengineering avoids false confidence.

The present data suggest that source construction is a first-order variable. MILimbEEG-only was a poor transport source, and aggressive balancing reduced performance by shrinking the source too far. At the same time, pooled raw and pooled dataset-balanced sources were broadly tied for the classical lead, even in matched-source comparisons. That combination matters. It argues against two common simplifications: first, that more healthy data is always better; second, that balancing alone will rescue transport. The more defensible interpretation is that source similarity and source composition matter, but no retrospective source recipe in the present public-data setting was strong enough to overcome the healthy-to-stroke gap.

Minimal adaptation also behaved in a revealing way. Ten-shot calibration substantially reduced ECE and shifted the operating point toward a more usable balance of sensitivity and specificity, yet AUROC changed very little. Ten-shot fine-tuning improved AUROC only modestly. The support-set resampling results sharpen this point further: AUROC was comparatively stable across support draws, whereas sensitivity and specificity varied markedly. In other words, the principal effect of limited subject-specific information in this benchmark was to help the model behave differently, not to make it rank much better. That is exactly the kind of distinction that matters if one imagines future lower-limb EEG control for rehabilitation or neuromodulation. A model that is weakly calibrated or threshold-fragile can be unusable even when AUROC appears acceptable.

The finding that classical baselines outperformed EEGNet is scientifically meaningful, not merely disappointing for deep learning. Under strong domain shift, small-sample target-domain adaptation, and imperfectly harmonised public EEG, simpler inductive biases may remain more stable than compact neural models. This does not imply that deep models are intrinsically unsuited to the problem. It does imply that, within this benchmark and tested model family set, additional retrospective architecture sweeps do not currently appear to be the decisive lever. The exploratory domain-generalisation branch supports the same conclusion: DG variants did not materially rescue transport.

The physiology audit is an equally important constraint. Canonical 16-channel performance was highest within dataset, but frontal and temporal control montages remained too competitive to support a strong motor-only interpretation. This paper should therefore not be read as evidence for a clinically deployable EEG biomarker of lower-limb motor intent. Rather, it should be read as evidence that the field still lacks sufficiently confound-audited transport validation. This distinction matters because EEG can support useful prediction without necessarily reflecting a brain-specific source, and translational neuroengineering should not conflate those possibilities.[18] Several limitations should be stated plainly. First, this was a retrospective public-dataset analysis rather than a prospective harmonised study. Second, cross-dataset label harmonisation was necessarily coarse; `effort_rest` was the most defensible shared variable, but not a guarantee of physiological equivalence across healthy execution, healthy imagery, and stroke motor imagery. An imagery-only sensitivity analysis would be valuable future work, but it was not part of the locked benchmark reported here. Third, the few-shot support and test sets were trial-disjoint within participant but not session/timepoint-disjoint within Stroke2025, so the current manuscript does not establish session-held-out adaptation. Fourth, absolute transport performance was modest, and therefore the paper should not be overread as a positive control-signal paper. Fifth, the physiology evidence was insufficient for a strong motor-specificity claim. Sixth, the public-data design did not permit live false-trigger endpoints or shown-output workflow assessment. That omission matters because biosignal-driven assistive interfaces can impose workload and usability burdens that are not captured by nominal control performance alone.[19, 20] None of these limitations negates the contribution. They define its scope: this is a translational constraint paper, not a deployment paper.

## Conclusions

Healthy-to-stroke transport of lower-limb EEG remained weak in this public-data benchmark. Within this benchmark, the next decisive gains are more likely to come from harmonised prospective validation than from further retrospective model iteration. Such validation should place healthy cohorts and cohorts involving participants with stroke under a locked task ontology, explicit EOG/EMG/kinematic confound channels, prespecified adaptation rules, and clinically anchored operating-point endpoints. The current manuscript provides the rationale for that next step.

## List of abbreviations

AUPRC: area under the precision-recall curve
AUROC: area under the receiver operating characteristic curve
BCI: brain-computer interface
CI: confidence interval
CORAL: correlation alignment
CSP: common spatial patterns
DANN: domain-adversarial neural network
DG: domain generalisation
ECE: expected calibration error
EEG: electroencephalography
EMG: electromyography
EOG: electrooculography
LDA: linear discriminant analysis
TSLR: tangent-space logistic regression
VREx: variance risk extrapolation

## Declarations

### Ethics approval and consent to participate

This study involved only secondary analysis of previously released public, de-identified human EEG datasets and no new participant recruitment, intervention, interaction, or identifiable-data handling by the authors. Ethics approval and participant consent for the original data collection were reported by the source-study investigators in the EEGMMIDB, MILimbEEG, and Stroke2025 publications [10–13]. No new human-participant data were collected by the authors for the present study, and no additional participant-consent procedures were undertaken for this secondary analysis.

### Consent for publication

Not applicable.

### Availability of data and materials

The datasets analysed during the current study are publicly available from EEGM- MIDB on PhysioNet (https://physionet.org/content/eegmmidb/; dataset DOI https://doi.org/10.13026/C28G6P) [10, 11, 21], from the MILimbEEG public dataset release on Mendeley Data (https://doi.org/10.17632/x8psbz3f6x.2) and its Data in Brief publication [12], and from the Stroke2025 dataset paper (https://doi.org/10.1038/s41597-025-04618-4) and linked figshare repository (https://doi.org/10.6084/m9.figshare.27130299) [13]. Aggregate harmonisation tables, source-sampling controls, and the supplementary figure PDF are included in this submission package and Additional file 1.

The analysis code used to generate the reported experiments and figures is publicly available in the lower-limb-eeg-transport repository (https://github.com/danielchoi0315/lower-limb-eeg-transport).[17] Dependencies and local dataset-directory requirements are documented in the public repository.

### Authors’ contributions

DC conceived the study, designed the benchmark, developed the software, performed the primary analyses, generated the figures, and drafted the manuscript. AC contributed to study design, methodology, software implementation, formal analysis, data curation, and manuscript revision. QL contributed to methodology, formal analysis, data curation, and manuscript revision. JP supervised the study, contributed to conceptualization and methodology, interpreted results, and critically revised the manuscript. All authors read and approved the final manuscript.

## Acknowledgements

The authors acknowledge the maintainers of EEGMMIDB, MILimbEEG, and Stroke2025 for making these public resources available.

## Additional file

### Additional file 1

File name: supplementary_sanity_control_and_matched_source_figures.pdf Format: PDF

Title: Supplementary sanity-control and matched-source figures Description: Contains Supplementary Figures S1 and S2 and their legends.

