## Additional file 1 for "A retrospective public external benchmark of healthy-to-stroke lower-limb EEG transport highlights source construction, adaptation burden, and confound sensitivity"

### Additional file 1: Supplementary sanity-control and matched-source figures

Junho Park[1*]

This file contains Supplementary Figures S1 and S2 and their legends.

### Supplementary Figure S1. Sanity controls.

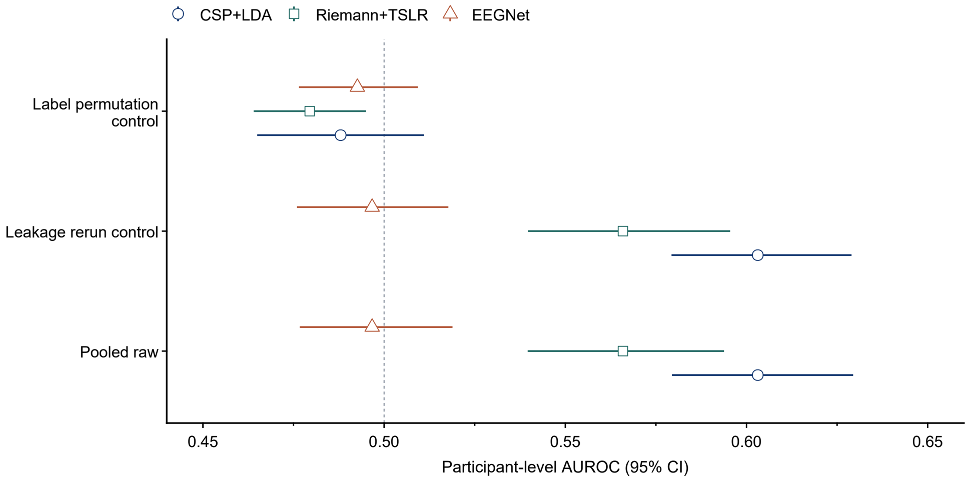


**Supplementary Figure S1. Sanity controls.** Label-permutation and leakage-control analyses for the leading pooled raw condition. Label permutation fell toward chance-like performance, whereas the leakage rerun reproduced the leading classical transport result.

### Supplementary Figure S2. Sample-size-matched source comparison.

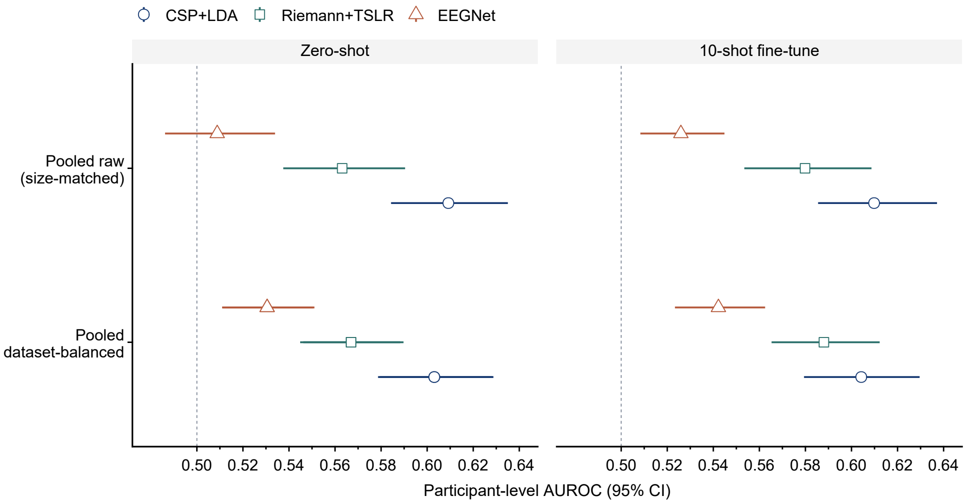


**Supplementary Figure S2. Sample-size-matched source comparison.** Zero-shot and 10-shot fine-tuned AUROC for pooled raw and pooled dataset-balanced source recipes under matched effective source size. The close alignment of the two recipes supports a restrained interpretation of source-balancing effects.
